# m6A RNA methylation facilitates pre-mRNA 3’-end formation and is essential for viability of *Toxoplasma gondii*

**DOI:** 10.1101/2021.01.29.428772

**Authors:** Michael J. Holmes, Leah R. Padgett, Matheus S. Bastos, William J. Sullivan

**Author notes:** Corresponding author: Indiana University School of Medicine, 635 Barnhill Drive, MS A418C, Indianapolis, IN 46202.

## Abstract

*Toxoplasma gondii* is an obligate intracellular parasite that can cause serious opportunistic disease in the immunocompromised or through congenital infection. To progress through its life cycle, *Toxoplasma* relies on multiple layers of gene regulation that includes an array of transcription and epigenetic factors. Over the last decade, the modification of mRNA has emerged as another important layer of gene regulation called epitranscriptomics. Here, we report that epitranscriptomics machinery exists in *Toxoplasma,* namely the methylation of adenosines (m6A) in mRNA transcripts. We identified novel components of the m6A methyltransferase complex and determined the distribution of m6A marks within the parasite transcriptome. m6A mapping revealed the modification to be preferentially located near the 3’-boundary of mRNAs within the consensus sequence, YGCAUGCR. Knockdown of the m6A writer enzyme METTL3 resulted in diminished m6A marks, loss of a target transcript, and a complete arrest of parasite replication. Furthermore, we examined the two proteins in *Toxoplasma* that possess YTH domains, which bind m6A marks, and showed them to be integral members of the cleavage and polyadenylation machinery that catalyzes the 3’-end processing of pre-mRNAs. Together, these findings establish that the m6A epitranscriptome is essential for parasite viability by contributing to the processing of mRNA 3’-ends.

**Author Summary:** *Toxoplasma gondii* is a parasite of medical importance that causes disease upon immuno-suppression. Uncovering essential pathways that the parasite uses for its basic biological processes may reveal opportunities for new anti-parasitic drug therapies. Here, we describe the machinery that *Toxoplasma* uses to modify specific adenosine residues within its messenger RNAs (mRNA) by N6-adenosine methylation (m6A). We discovered that m6A mRNA methylation is prevalent in multiple stages of the parasite life cycle and is required for parasite replication. We also establish that m6A plays a major role in the proper maturation of mRNA. Two proteins that bind m6A modifications on mRNA associate with factors responsible for the cleavage and final processing steps of mRNA maturation. Since all of the machinery is conserved from plants to *Toxoplasma* and other related parasites, we propose that this system operates similarly in these organisms.

## Introduction

*Toxoplasma gondii* is an obligate intracellular parasite that accounts for nearly a quarter of lethal foodborne illnesses in the U.S.A. (1). *Toxoplasma*-induced illnesses are estimated to cost the U.S. economy three billion dollars annually (2). In addition, toxoplasmosis is a major opportunistic infection in immune compromised patients, such as those with HIV/AIDS, and can cause congenital birth defects (3). Due to its ubiquitous distribution around the globe (4) coupled with the average seroprevalence estimated to be approximately a third of human population worldwide (3), *Toxoplasma* constitutes a major health burden for which relatively few therapeutic options exist.

*Toxoplasma* can infect any warm-blooded animal and is usually acquired by ingesting one of two encysted forms: oocysts expelled by feline hosts into the environment or tissue cysts in undercooked animal products (3). Once ingested, the parasites pass through the rapidly growing tachyzoite phase to disseminate throughout the host, then convert into quiescent bradyzoites within tissue cysts (3). Tissue cysts persist in various organs, particularly the brain, leading to a life-long infection that is currently incurable (3). If host immunity wanes, bradyzoites can reconvert into replicative tachyzoites, causing localized necrosis and potentially death (3). Frontline chemotherapeutics target processes critical for replication, such as pyrimidine biosynthesis and translation within the parasite’s plastid-like apicoplast and have little to no impact on tissue cysts (5). These limited drug treatments suffer significant drawbacks that include serious adverse effects, potential allergy, and differing efficacy between parasite strains (5). As such, there is a need to discover essential processes in the parasite that have biological distinctions from the host in order to identify novel targets that could be leveraged for new anti-*Toxoplasma* therapies.

Over the course of the last decade, mRNA methylation has emerged as an important new layer of eukaryotic gene regulation (6-11). The most abundant internal mRNA modification is the N6-methylation of adenosines (m6A) (12). Due to the analogies that methylation of internal mRNA adenosines share with the reversible modification of DNA residues (termed the epigenome), the content and dynamic profile of methylated mRNA has been dubbed the epitranscriptome. Similarly, the regulatory machinery that deposits, confers function, and removes the methyl marks are referred to as m6A ‘writers’, ‘readers’, and ‘erasers’, respectively (9).

The m6A mark has been implicated in nearly all facets of mRNA biology, ranging from playing a role in alternative splicing and nuclear export to influencing transcript localization, stability, translation, and turnover (8, 9, 12, 13). Consequently, the m6A mark and the proteins that regulate and recognize it are instrumental in diverse biological processes including cell differentiation, meiosis, sex determination, developmental patterning, and stress responses, as well as disease states in various organisms (6-8, 12, 13).

Across eukaryotes, the mRNA methyltransferase writer complex is comprised of the three conserved subunits METTL3, METTL14, and WTAP, as well as various accessory proteins that are not conserved across species (6, 7, 12). While the m6A mark itself may alter local secondary structures by lessening the strength between A:U base pairs in mRNAs (14, 15), the majority of m6A-directed activities stem from the direct recognition of the methylated nucleotide by RNA-binding proteins, termed m6A readers. The best characterized readers are YTH domain-containing proteins, though additional proteins have also been demonstrated to recognize the m6A mark (7-9, 12, 13). Selective removal of the m6A mark occurs through the erasers, Fe(II)/α-ketoglutarate-dependent dioxygenases in the protein ALKBH family (7, 9, 12).

Phylogenetic evidence indicates that the m6A epitranscriptome system is present in each of the eukaryotic kingdoms, including many animals, plants, fungi, and the Alveolate protozoa, including *Plasmodium spp.* and *Toxoplasma* (8). Here, we systematically investigate each identifiable member of the m6A machinery in *Toxoplasma* and survey the transcriptome-wide prevalence of m6A in tachyzoites and in parasites undergoing bradyzoite formation. We determine that writer functionality is essential for parasite viability, identify a parasite-specific member of the writer complex, discover that the m6A mark is localized in close proximity to the 3’-end of transcripts, and demonstrate that the YTH-family reader proteins constitute part of the machinery involved in processing the cleavage and polyadenylation of transcripts at their 3’-termini. Together, these findings establish that m6A is a critical mRNA modification that is widespread across multiple stages of the *Toxoplasma* life cycle and relies on parasite-specific components that may serve as attractive drug targets.

## Results

### The m6A mark is present throughout asexual *Toxoplasma* stages

To determine whether m6A is present in *Toxoplasma,* we first performed immunofluorescent assays using an anti-m6A antibody. Examination of extracellular tachyzoites (ME49 strain) revealed an enriched signal in the cytoplasm that appeared in a slightly punctate pattern (Fig. 1A, panel i). A corresponding no antibody control, which was performed at the same time, indicated that the staining was specific (Fig. 1A, panel ii). We also examined intracellular tachyzoites and bradyzoites generated by treating parasites with alkaline media for 6 days. The m6A mark was primarily localized to the cytoplasm in both of these life stages (Fig. 1A, panels iii and iv). An anti-m6A dot blot performed on total RNA and genomic DNA extracted from purified tachyzoites revealed an enrichment of m6A in parasite RNA (Fig. 1B). The quality of the parasite purification was assessed by the lack of a detectable host-specific 28S rRNA band in the RNA sample, indicating that very little to no contaminating host nucleic acids were present in the sample (data not shown). Together, these data demonstrate that m6A is a feature of *Toxoplasma* RNA.

**Figure 1.**
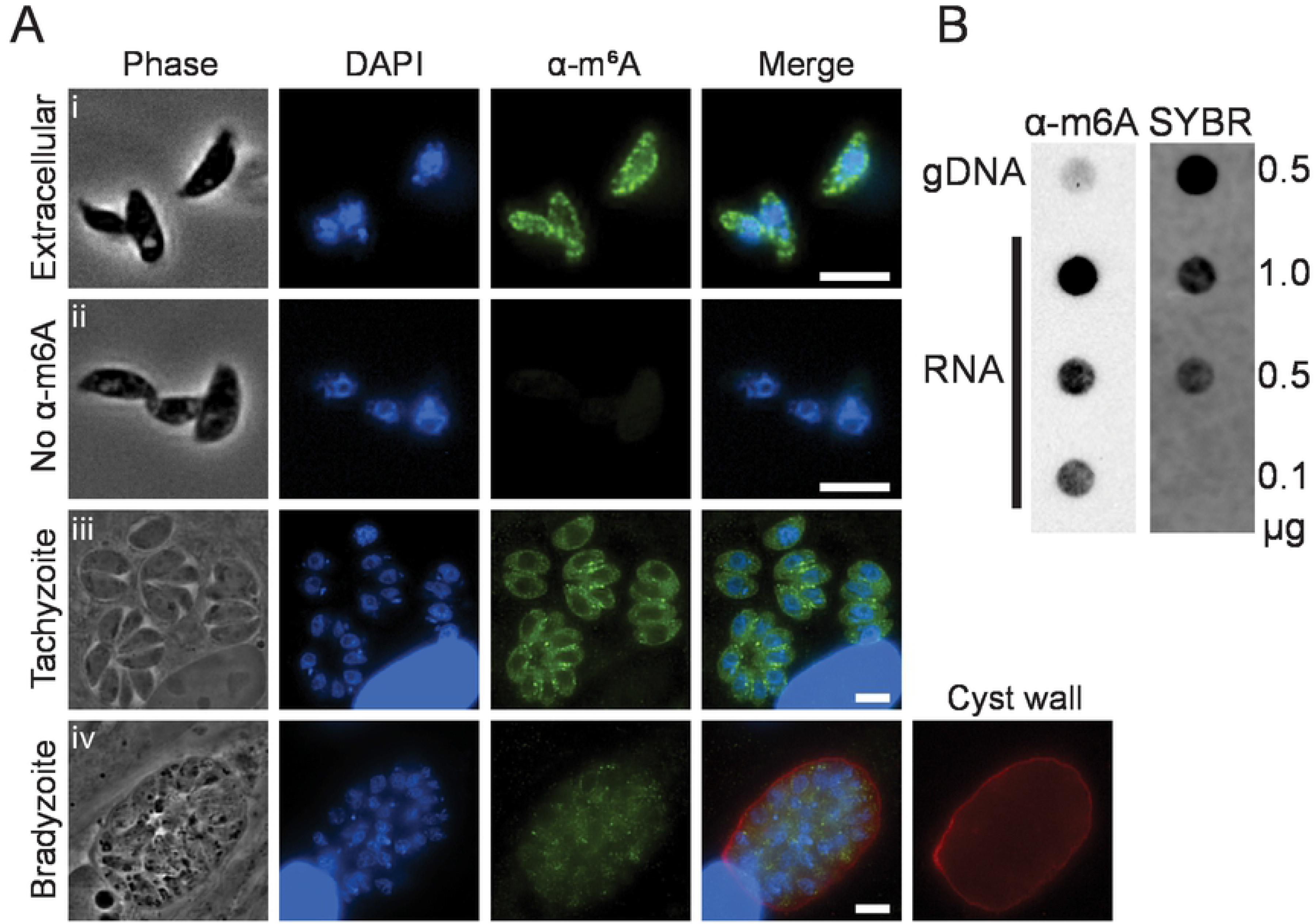
*Toxoplasma* RNA is modified with the m6A mark. A) Immunofluorescence assays show m6A-specific marks enriched in the cytoplasm of i) extracellular tachyzoites compared to ii) a control without primary antibody. The m6A mark is also present in the cytoplasm of iii) intracellular tachyzoites and iv) stress-induced bradyzoites (cyst wall is stained with DBA). Scale bars represent 5 μm. B) *Toxoplasma* genomic DNA and differing amounts of RNA, as indicated, were immunoblotted for the m6A mark. SYBR Gold staining serves as a loading control.

### Purification of the *Toxoplasma* m6A writer complex reveals unique features

The components of the m6A writer complex, which consists of the catalytic METTL3, the catalytically inactive METTL14, and the associated protein WTAP are conserved in *Toxoplasma* based on BLASTP homology searches (Table 1). It should be noted that WTAP is currently annotated as two separate genes on ToxoDB, but it is likely to be a single transcript (see next section). Of importance, each of three of the *Toxoplasma* m6A writer orthologues are critical for parasite fitness according to a genome-wide CRISPR screen (Table 1) (16), suggesting that adenosine methylation of mRNA is important and required for tachyzoite growth and replication.

**Table 1.** m6A writer components in *Toxoplasma*.

| Gene ID | Description | Gene Name | CRISPR score |
| --- | --- | --- | --- |
| Writers |  |  |  |
| TGGT1_217350 | putative methyltransferase MTA70 | METTL3 | -4.05±1.62 |
| TGGT1_268840 | putative N6-adenosine-methyltransferase | METTL14 | -4.25±0.88 |
| TGGT1_212960A<sup>a</sup> | hypothetical protein | WTAP | -4.87±0.79 |
| TGGT1_212960B<sup>a</sup> | hypothetical protein | WTAP | -3.59±0.38 |
| Readers |  |  |  |
| TGGT1_201200 | zinc finger (CCCH type) motif-containing protein | YTH1 | -3.11±0.61 |
| TGGT1_204070 | YT521-B family protein | YTH2 | -4.28±1.13 |
| Erasers |  |  |  |
| None detected |  |  |  |

<sup>a</sup> WTAP is misannotated as two genes

We tagged each of identified m6A writer proteins at their endogenous C-termini with an HA epitope in RH strain parasites lacking *KU80* and *HXGPRT* genes (RHΔΔ) using single crossover recombination (17, 18). Consistent with observations from other eukaryotes and with their proposed role in mediating co-transcriptional mRNA methylation (9), each of the writer proteins were predominantly localized in the parasite nuclei (Fig. 2A). Whereas METTL3^HA^ and WTAP^HA^ were easily detected, the METTL14^HA^ signal was comparatively faint and required more sensitive exposure settings for adequate visualization. A western blot of each tagged line revealed a single band of the expected size; the data also support that WTAP is misannotated on ToxoDB and should be a single gene (Fig. 2B). We also note that while WTAP and METTL3 proteins are expressed in similar amounts, METTL14 is significantly expressed at substoichiometric levels, consistent with the diminished detection in the immunofluorescence microscopy (Fig. 2B-C). Whether this result is indicative of a potential regulatory function for METTL14 as a limiting factor for the writer complex remains to be determined.

**Figure 2.**
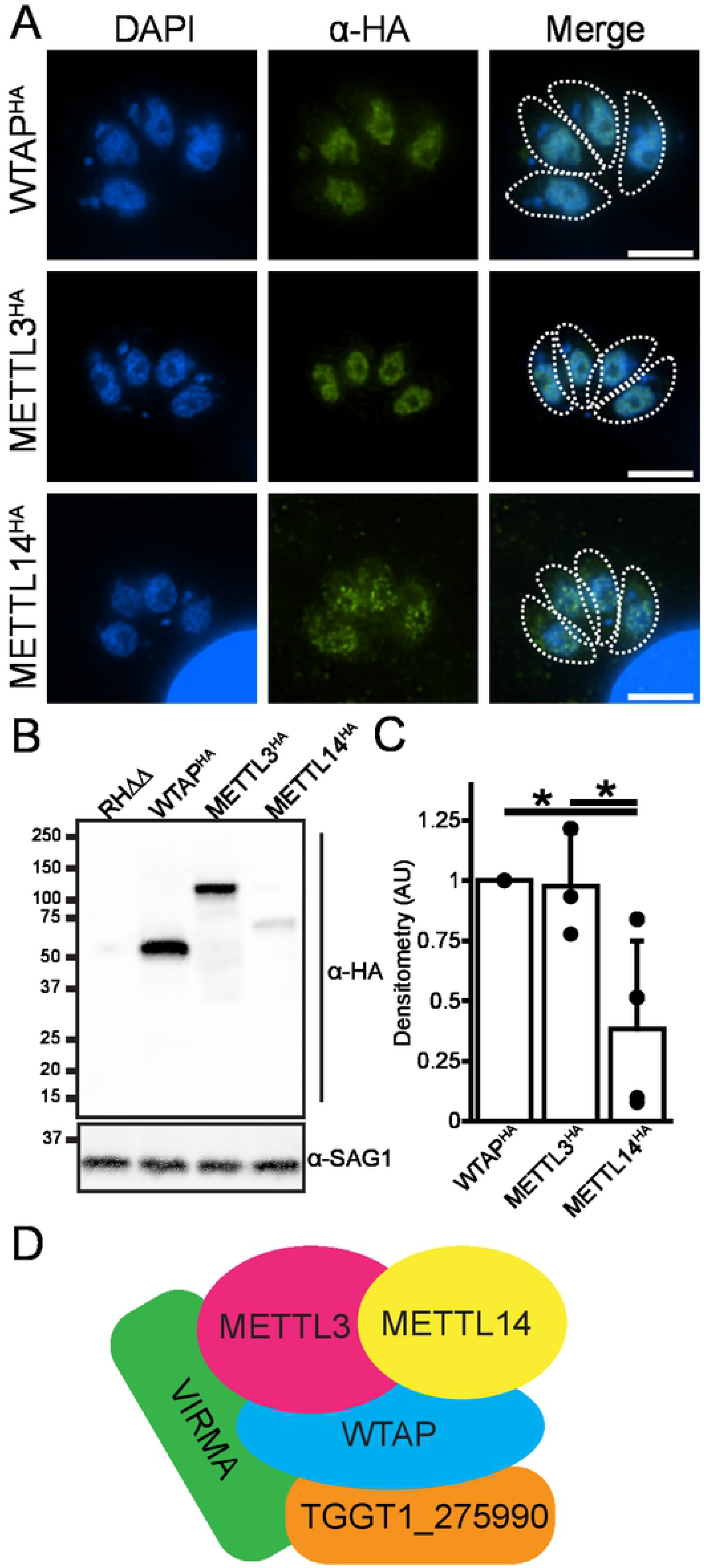
Components of *Toxoplasma* m6A writer complex. A) Immunofluorescence assay of endogenously HA-tagged WTAP, METTL3, and METTL14 subunits of the writer complex. Each protein is localized to the parasites’ nuclei. The outlines of parasites are shown in the merge panel for reference. B) Western blot of endogenously HA-tagged writer complex members. C) Relative quantification of writer complex member expression as assessed by Western blot densitometry. Data are presented as average ± standard deviation normalized to WTAP^HA^ signal. * represents p ≤ 0.05 as assessed by Student’s t-test assuming unequal variances. D) Schematic summarizing the *Toxoplasma* writer complex membership as identified by co-immunoprecipitation experiments outlined in Table 2.

METTL3, METTL14, and WTAP are widely conserved amongst eukaryotes as a “core” complex that delivers the m6A mark, but accessory proteins involved in this writer complex can differ among species. There are no apparent orthologues of any accessory proteins from other organisms in *Toxoplasma* based on sequence homology. To validate the associations between *Toxoplasma* METTL3^HA^, METTL14^HA^, and WTAP^HA^, and discover possible accessory writer proteins, we immunoprecipitated each of the HA-tagged proteins from nuclear extracts of freshly lysed tachyzoites and identified copurifying proteins by mass spectrometry. Both METTL3^HA^ and METTL14^HA^ were robustly pulled down together in every experiment (Table 2). Interestingly, although WTAP was not significantly enriched in either METTL3^HA^ or METTL14^HA^ pulldowns, a few peptides were present in one experiment of each (Table 2, Table S1).

**Table 2.** Interactors of the m6A writer complex. The m6A writer complex was identified from immunoprecipitation of HA-tagged METTL3, METTL14, and WTAP. Peptide counts from two independent experiments are shown.

| Gene ID | Description | METTL3<sup>HAa</sup> | METTL14<sup>HAa</sup> | WTAP<sup>HAa</sup> |
| --- | --- | --- | --- | --- |
| TGGT1_217350 | putative methyltransferase MTA70 (METTL3) | **36\|198<sup>c</sup>** | **103\|100** | 0\|0 |
| TGGT1_268840 | putative N6-adenosine-methyltransferase (METTL14) | **32\|143** | **81\|64** | 0\|0 |
| TGGT1_212960A<sup>b</sup> | hypothetical protein (WTAP) | 0\|0 | 0\|0 | 0\|10 |
| TGGT1_212960B | hypothetical protein (WTAP) | 0\|5 | 0\|1 | **36\|159** |
| TGGT1_226660 | hypothetical protein (VIRMA) | 0\|1 | 0\|0 | **222\|710** |
| TGGT1_275990 | hypothetical protein | 0\|0 | 0\|0 | **18\|126** |

<sup>a</sup> protein that was immunoprecipitated
<sup>b</sup> misannotated WTAP fragment was manually included in this list
<sup>c</sup> bolded peptide counts are significantly enriched over parental ( $P_{adj} \leq 0.05$ ; Fisher's exact test with Benjamini-Hochberg correction)

In reciprocal pull down experiments, neither METTL3 nor METTL14 were found in WTAP^HA^ immunoprecipitates. Rather, WTAP associated with two hypothetical proteins, TGGT1_226660 and TGGT1_275990, with high significance. Although neither protein possesses any annotated domains, homology searches conducted on the HHPRED webserver (19) revealed a region in the N-terminus of TGGT1_226660 that displayed homology with the PF15912 domain corresponding to the N-terminal domain of VIRMA, a well-documented component of the m6A writer complex that helps guide the writer complex to the stop codon and 3’-UTR in higher eukaryotes (6, 20). Interestingly, when performing orthology searches of TGGT1_275990, we discovered that it is restricted to coccidian parasites, suggesting that this m6A writer-associated protein may be a unique feature present in select apicomplexan parasites.

### The m6A epitranscriptome in tachyzoites and early bradyzoites

In higher eukaryotes, deposition of m6A has been shown to be dynamic and required for the efficient progression of developmental and biological processes (6, 8, 12). We therefore characterized the m6A epitranscriptome as *Toxoplasma* converts from tachyzoites to early bradyzoites. We performed m6A-enriched RNA sequencing using the MeRIPseq method (21) in parallel with RNAseq in ME49 tachyzoites grown under normal conditions and under alkaline pH for 24 hours, a stress commonly used to induce bradyzoite conversion *in vitro* (22). RNAseq measurements from parasites subjected to this induction trigger showed an upregulation of 295 gene transcripts, including those encoding proteins associated with bradyzoites, such as the cyst wall antigens CST1 and MAG1, the stage-specific enzymes EN01 and LDH2, and AP2 transcription factors associated with early bradyzoite development. Additionally, among the 19 downregulated genes were the canonical tachyzoite surface antigen SAG1 (23, 24) (Fig. 3A, Table S2). While it is important to underscore that these parasites are not mature bradyzoites, our results are consistent with the gene expression pattern expected of tachyzoites that have begun the conversion into bradyzoites.

**Figure 3.**
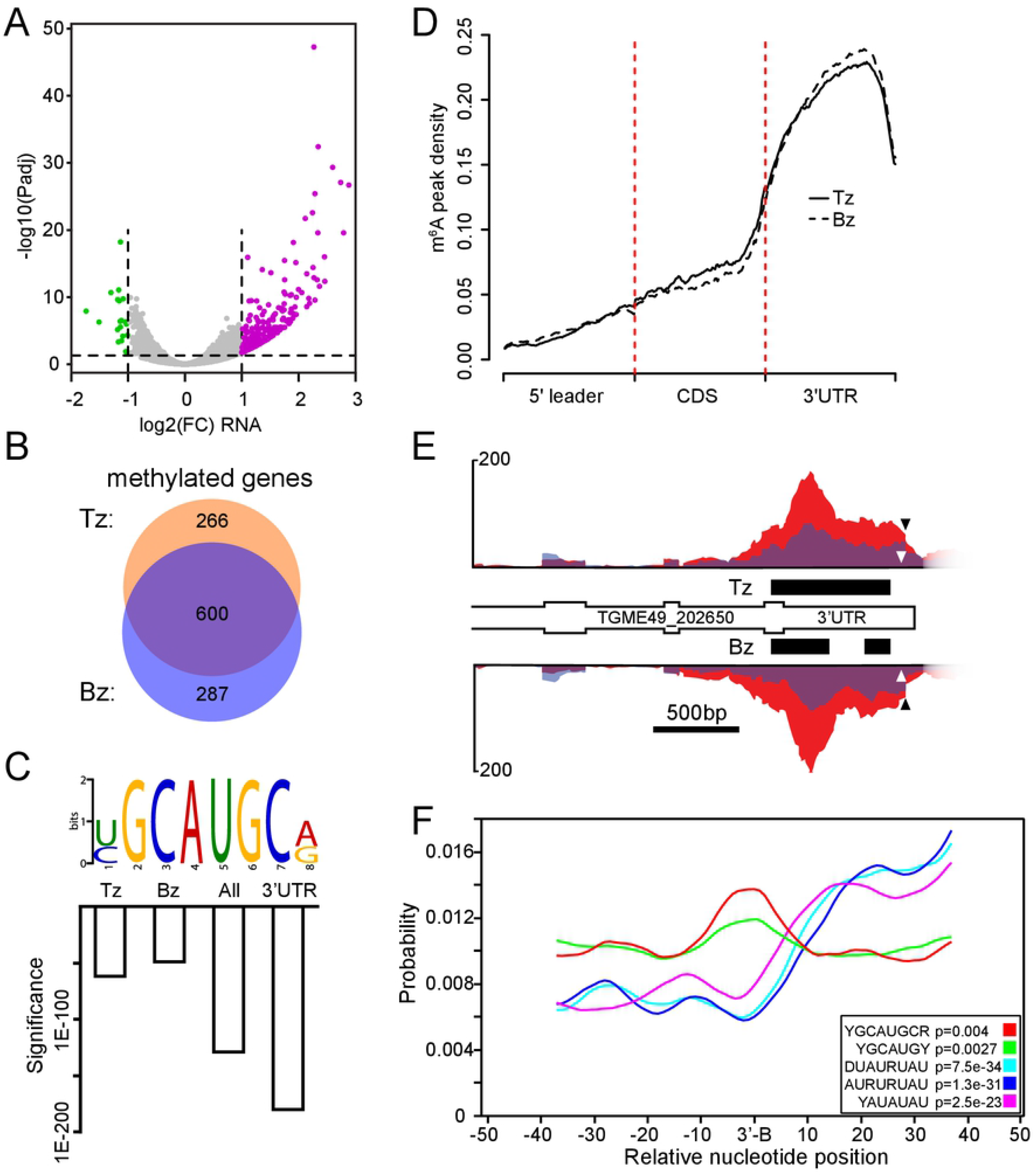
The m6A mark is enriched near mRNA 3’-boundaries. A) Volcano plot displaying gene expression changes upon 24 h treatment with alkaline stress. The 295 upregulated genes, displayed in purple, include classic bradyzoite markers and the 19 downregulated genes are displayed in green. B) Venn diagram showing the overlap between m6A-modified genes from tachyzoites (Tz) and parasites after 24 h stress treatment (Bz). C) Motif identification of m6A-enriched sequencing. Significance (e value) is displayed in a bar chart for sequencing data from tachyzoites (Tz), bradyzoite-induced sample (Bz) and a combination of all m6A-enriched sequences (All). The same motif was independently identified from all annotated 3’-UTRs in ToxoDB. D) Metagene plot showing average distribution of m6A density along a normalized transcript from a single tachyzoite and bradyzoite-induced replicate. The m6A mark is most abundant in annotated 3’-UTRs. All three replicates from each condition displayed similar distributions. E) Distribution of m6A-enriched MeRIPseq (red) and input RNAseq (blue) reads from tachyzoite (Tz) and alkaline stressed (Bz) samples. m6A-containing peaks are displayed as black bars for both conditions. The m6A motif is indicated by the white arrowhead and the putative 3’-boundary is shown by the black arrowhead. All data are shown to scale. F) m6A-like motifs are enriched near mRNA 3’-boundaries (3’-B) in *Toxoplasma* transcripts. Each motif variant detected in this study is displayed. An enrichment of AU-rich elements was also seen downstream of the 3’-boundary. Statistical significance of the localized enrichment (Bonferroni-corrected one-tailed binomial test) is shown on the bottom right of the panel for each individual motif.

To analyze our datasets for m6A-containing transcripts, we utilized the MeTPeak software package that has been tailored for detecting m6A-enriched regions in MeRIPseq experiments (25). We detected 1,896 peaks spread across 866 genes in our tachyzoite dataset and 1,841 peaks amongst 687 genes during early bradyzoite formation (Table S3). The majority of the m6A-containing transcripts are shared between the two conditions (Fig. 3B). We queried the MeRIPseq datasets for gene ontology enrichment of biological processes and molecular functions, however no terms were significantly enriched. We then used MeTDiff (26), which takes the MeRIPseq and RNAseq data from all conditions into account to call m6A-containing peaks, to assess differential m6A methylation between the two conditions. This process detected a total of 6,777 peaks spread across 3,272 genes, however none were differentially methylated after this 24 hour stress-induction period (Table S3). These results show that m6A is an abundant modification on mRNAs encoding factors of diverse functions in both tachyzoites and early bradyzoites.

### m6A marks are enriched near the 3’-boundary of mRNAs

To determine whether the modified mRNAs contain a consensus sequence targeted for m6A methylation, we utilized the DREME tool (27) which revealed a highly enriched motif: YGCAUGCR, in which Y represents a pYrimidine and R represents a puRine. The YGCAUGCR motif was identified in both tachyzoites and early differentiating bradyzoites, as well as in a combination of all the peaks identified by MeTDiff (Fig. 3C, Fig. S1). This motif contains a centrally-located adenosine that would represent the methylated residue. Variations on a second AU-rich motif were also reliably identified, though with less significance (discussed below, Fig. S1).

In other organisms, the distribution of m6A marks within the transcriptome are known to be enriched in specific transcript regions. For example, m6A marks tend to localize near the stop codon in mammals (8), within the CDS in *Plasmodium falciparum* (28), and within the polyA tail in *Trypanosoma brucei* (29). A metagene plot of m6A peaks demonstrates that they are highly enriched in *Toxoplasma* 3’-UTRs (Fig. 3D). A specific example of a methylated transcript, the hypothetical protein TGME49_202650, shows increased MeRIPseq signal within its 3’-UTR and has m6A peaks (denoted as black bars) in both tachyzoite and early bradyzoite samples (Fig. 3E). When we examined the annotated peaks of this gene for the presence of the presumptive m6A YGCAUGCR motif, we found it located just outside the boundary of the called peaks near the 3’-end of the transcript (Fig. 3E, white arrowhead). We also noted that the YGCAUGCR motif was located only 22nt upstream from a sharp decline in the mapping of the RNAseq data (Fig. 3E, black arrowhead) suggestive of the 3’-boundary of the transcript.

We suspected that the m6A motif may be systemically enriched near the 3’-boundary of mRNAs but that m6A-enriched regions may be difficult to detect due to the sharp decline in sequencing reads near the polyadenylation site. To assess this, we used the DREME algorithm to unbiasedly identify motifs that were enriched within the 3’-UTRs. We limited our analysis to the 6109 protein coding genes with 3’-UTRs that were least 9 nt long since the YGCAUGCGR motif is 8 nt long. The top identified motif was a variation on the AU-rich motif previously identified in our MeRIPseq dataset, whereas the second was the YGCAUGCR motif that we propose is an m6A consensus sequence (Fig. 3C, Fig. S1). We used the CentriMo algorithm (30) to determine whether these motifs were enriched near the annotated 3’-boundaries from the same gene set (Fig. S1). Whereas the AU-rich motifs were located downstream, the m6A motif was significantly enriched in a region closely associated with the annotated 3’-boundary of mRNAs (Fig. 3F). This is strongly indicative that at least a subset of *Toxoplasma* transcripts is modified with the m6A mark near their 3’-boundary.

### METTL3 depletion reduces steady state level of mRNA with m6A motif and arrests parasite replication

Since our analyses indicated that a substantial subset of m6A marks are present at the 3’-end of transcripts, we wanted to assess whether disruption of the m6A system would impact mRNAs that are marked near their 3’-boundary. In order to verify this, we endogenously tagged the catalytic subunit of the writer complex, METTL3, at its C-terminus with an HA epitope and auxin-inducible degron (AID) tag in RH strain parasites engineered to contain TIR1; this allows rapid degradation of the AID-fusion protein upon the addition of indole-3-acetic acid (IAA) (31). Western blot analysis revealed near complete loss of the METTL3^HA-AID^ protein in as little as 15 minutes following addition of 500 μM IAA in the culture medium (Fig. 4A). We also noted a decrease in SAG1 expression after extended periods of IAA incubation, possibly due to decreased parasite viability following extended loss of METTL3 (see below). Consistent with its role as the catalytically active m6A methyltransferase, depletion of METTL3 for 4 h diminished m6A levels in the parasite (Fig. 4B).

**Figure 4.**
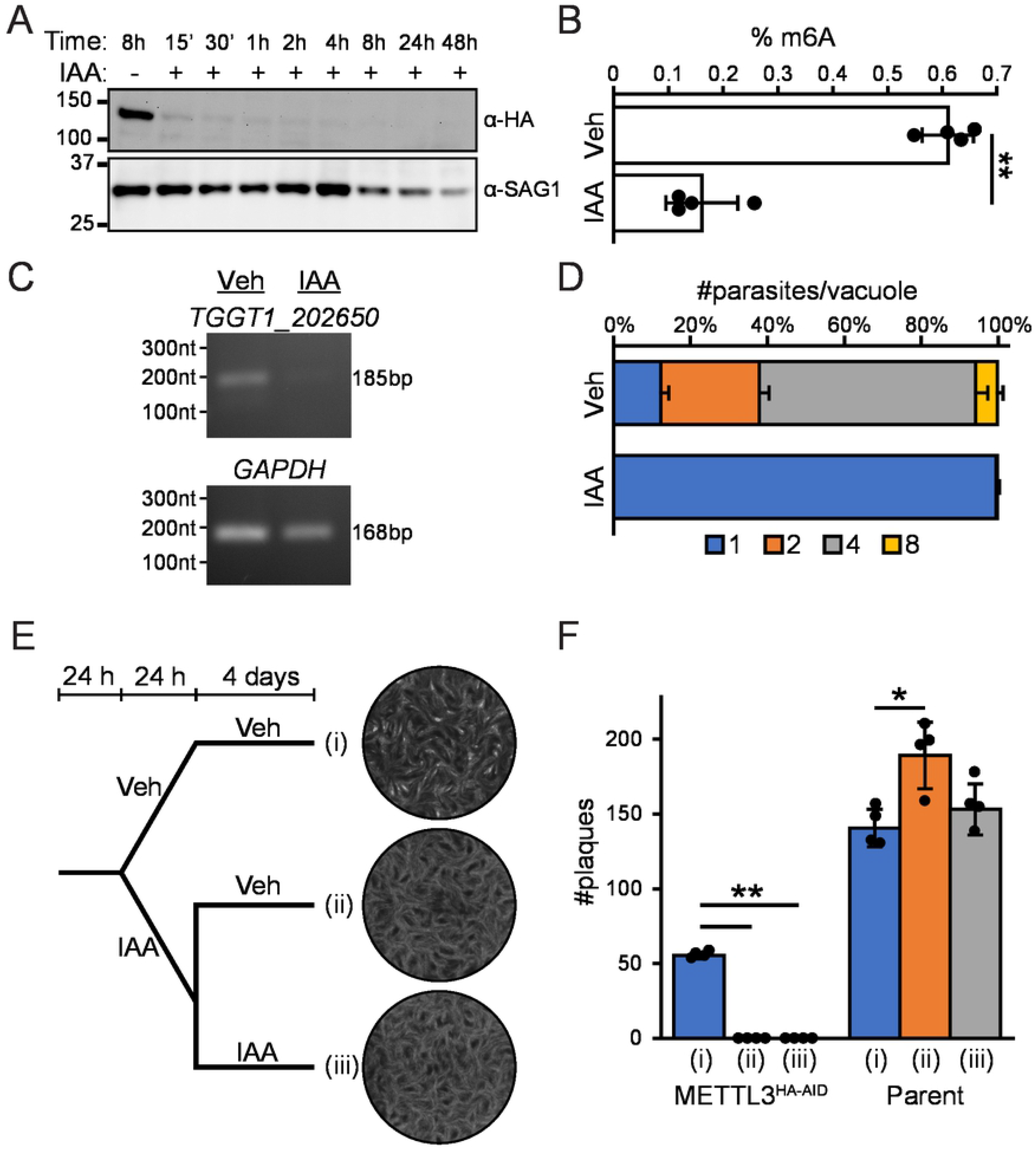
Effect of METTL3 depletion in *Toxoplasma*. A) Western blot of METTL3^HA-AID^ parasites treated with 500 μM IAA or DMSO vehicle for the indicated times. Rapid depletion of METTL3^HA-AID^ is seen upon addition of IAA. A decreased signal in the SAG1 loading control is also apparent upon prolonged IAA exposure. B) Quantitation of m6A levels after treatment of METTL3^HA-AID^ parasites with 500 μM IAA for 4 h. C) RT-PCR analysis of an m6A-motif containing transcript, *TGGT1_202650,* and a control mRNA, *GAPDH,* which does not encode the motif near its 3’-boundary. D) Replication assay of METTL3^HA-AID^ parasites upon treatment with 500 μM IAA or DMSO vehicle for 16 h. Data are presented as the average of three replicates ± standard deviation. E) Plaque assay of METTL3^HA-AID^ parasites and schematic representation of IAA dosing regimens. Parasites were allowed to grow for 24 h before undergoing one of three treatment regimens. Parasites were treated with i) DMSO vehicle for 24 h followed by 4 days in fresh media with DMSO; ii) 500 μM IAA for 24 h followed by 4 days in fresh media with DMSO; or iii) 500 μM IAA followed by 4 days in fresh media with IAA. Representative images of each treatment regimen for METTL3^HA-AID^ parasites are displayed. F) Quantitation of the plaque assay described in (E) for the METTL3^HA-AID^ and the RH-TIR1 parental strains. Values are presented as the average of 4 replicates ± standard deviation. Statistical significance was assessed by Student’s t-test assuming unequal variances. * represents p ≤ 0.05, ** represents p ≤ 0.001. All experiments were repeated at least twice with similar results.

In order to assess how METTL3 depletion impacted an m6A-marked transcript, we isolated total RNA from METTL3^HA-AID^ parasites that were treated with DMSO or IAA-treated for 4 h and performed an RT-PCR (Fig. 4C). *TGGT1_202650,* whose ME49 homologue is marked with the m6A motif near its 3’-boundary, was barely detectable following IAA treatment. In contrast, IAA had little impact on *GAPDH* mRNA levels, which lacks the m6A motif near its 3’-boundary. This finding suggests that failure to deliver m6A marks to the 3’-end of target mRNAs leads to loss of the transcript.

We next examined the effect of METTL3 depletion on parasite replication using a standard parasite counting assay. Infected HFF cells were incubated for 16 h in IAA or DMSO vehicle; parasites depleted of METTL3 exhibited arrested replication (Fig. 4D). In order to determine whether parasites can recover from a transient loss of METTL3 expression, we conducted modified plaque assays that used three different IAA treatment regimens (Fig. 4E). After an initial 24 h of growth, METTL3^HA-AID^ parasites were treated with IAA for 24 hours, at which point the media was replaced with or without IAA. There were no detectable plaques regardless of whether parasites were treated with a single pulse of IAA for 24 h or a sustained dose of IAA (Fig. 4E-F). These results indicate that parasites could not recover after depletion of m6A for 24 h. It is noted that IAA did not inhibit parasite plaquing of the parental line; however, the tagging of METTL3 moderately compromises plaquing efficiency compared to the parental line (Fig. 4F). Together, these findings highlight the importance of METTL3 and m6A for *Toxoplasma* viability and suggest a linkage between m6A and mRNA processing.

### YTH m6A reader proteins are core constituents of the cleavage and polyadenylation machinery in *Toxoplasma*

The m6A mark has been linked to a variety of biological processes, and the functional consequence of the modification is carried out by m6A reader proteins, most notably those containing the YTH RNA-binding domain (7-9). The *Toxoplasma* genome encodes two such proteins, TGGT1_201200 and TGGT1_204070, which we termed YTH1 and YTH2, respectively (Table 1). To investigate their role in *Toxoplasma,* we endogenously tagged each at their C-termini with an HA epitope in RHΔΔ parasites (17, 18). Western blot analysis of YTH1^HA^ and YTH2^HA^ with anti-HA revealed primary bands of predicted sizes at ~70 kDa and 60 kDa, respectively (Fig. 5A-B). Multiple lower molecular weight bands were consistently detected for YTH1^HA^ and may either indicate that this protein is proteolytically processed or particularly heat labile. Additionally, a minor higher molecular weight band was consistently detected for YTH2^HA^, suggestive of a post-translational modification(s). As seen for the m6A writer complex members, both YTH1^HA^ and YTH2^HA^ localize to the parasite nucleus (Fig. 5C).

**Figure 5.**
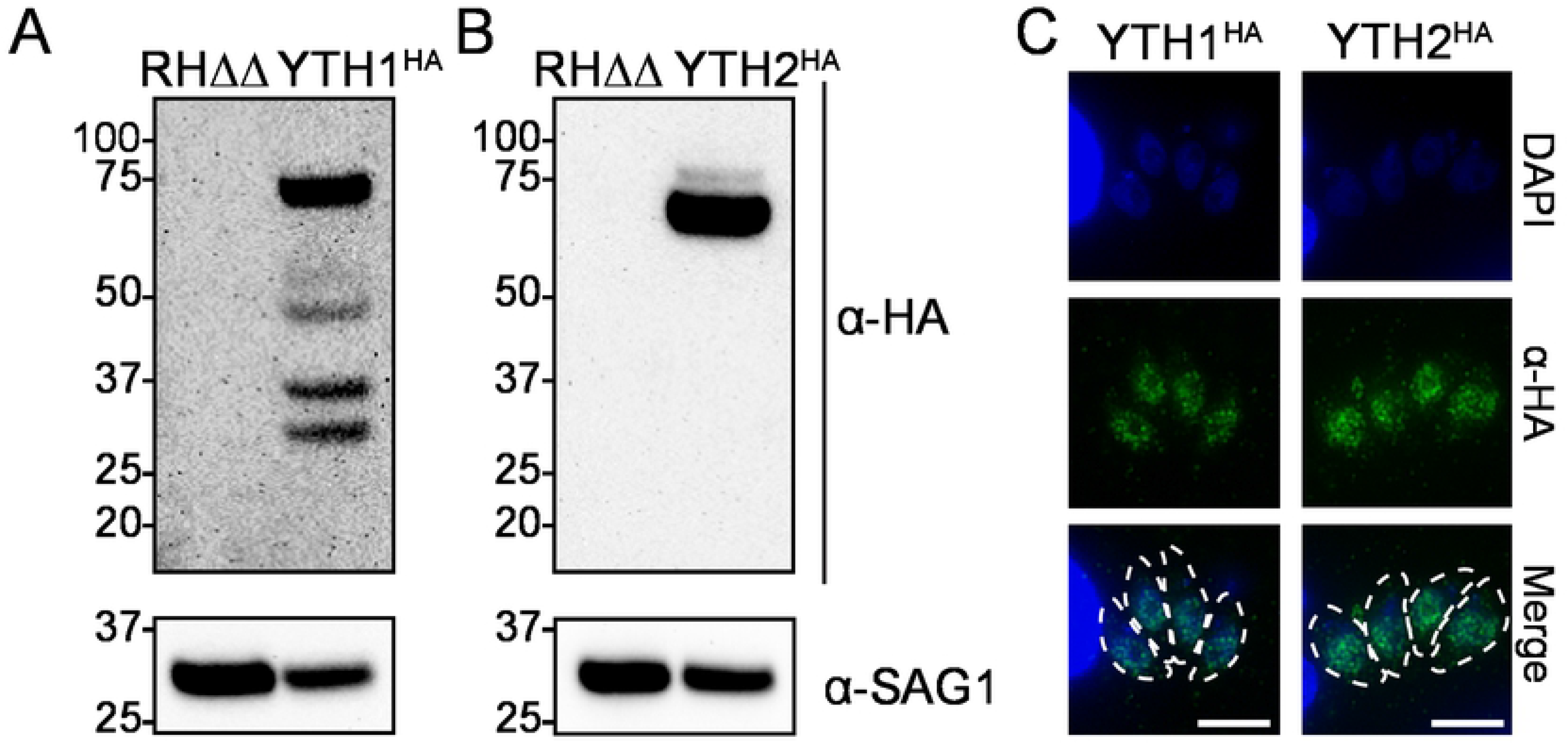
Localization of YTH domain-containing m6A reader proteins. A-B) Western blot of endogenously HA-tagged A) YTH1 and B) YTH2 reader proteins. Both proteins display reproducible additional bands that are not present in the RHΔΔ parental strain. C) Immunofluorescence assay of each HA-tagged YTH reader proteins. Outlines of the parasites are shown in the merge panel for reference. Scale bars represent 5 μm.

To gain insight into their cellular function, we immunoprecipitated each YTH protein from intracellular tachyzoites. Numerous proteins that co-purified with YTH1^HA^ are annotated in ToxoDB as core constituents of the 3’ cleavage and polyadenylation machinery (Table 3, Table S4), which coordinate the 3’-end formation of mRNAs. Additional interacting proteins include other putative RNA-binding proteins, many of which share clear sequence homology with other components of the 3’-end formation machinery (Table 3). Notably, the *Toxoplasma* YTH1 protein, which encodes N-terminal CCCH zinc finger (ZnF) domains and a C-terminal YTH domain, shares the same protein domain architecture and high homology with CPSF30 (Cleavage and Polyadenylation Specificity Factor 30) found in plants (32, 33).

**Table 3.** Proteins interacting with the YTH-domain m6A reader proteins. Interacting proteins were identified from immunoprecipitation of HA-tagged YTH1 and YTH2. Peptide counts from two biological replicates are shown.

| Gene ID | Product Description | Orthologue | YTH1<sup>HAa</sup> | YTH2<sup>HAa</sup> |
| --- | --- | --- | --- | --- |
| m6A-readers |  |  |  |  |
| TGGT1_201200 | Zinc finger (CCCH type) motif-containing protein | CPSF30 | **5\|7<sup>b</sup>** | 0\|0 |
| TGGT1_204070 | YTH domain-containing protein | CPSF30 | 0\|0 | **6\|6** |
| mRNA cleavage and polyadenylation factors |  |  |  |  |
| TGGT1_219440 | Cleavage and polyadenylation specificity factor subunit 2 | CPSF100 | **23\|15** | 0\|0 |
| TGGT1_224280 | CPSF A domain-containing protein | CPSF160 | **21\|29** | 0\|0 |
| TGGT1_230010 | Suf domain-containing protein | CSTF77 | **19\|19** | 0\|0 |
| TGGT1_247330 | Symplekin C domain-containing protein | Symplekin | **15\|12** | 0\|0 |
| TGGT1_294700 | Polynucleotide 5'-hydroxyl-kinase | CLP1 | **6\|8** | 0\|0 |
| TGGT1_254210 | RRM domain-containing protein | CSTF64 | **6\|6** | 0\|0 |
| TGGT1_268250 | WD repeats region domain-containing protein | WDR33 | **5\|6** | 0\|0 |
| TGGT1_285200 | Cleavage and polyadenylation specificity factor protein | CPSF73 | 4\|10<sup>c</sup> | 0\|0 |
| TGGT1_283740 | RRM domain-containing protein | CSTF64 | **4\|4** | 0\|0 |
| TGGT1_230230 | FIP1 domain-containing protein | FIP1 | **3\|3** | 0\|0 |
| TGGT1_268900 | RING-type domain-containing protein | RBBP6 | 0\|0 | **6\|8** |
| TGGT1_221190 | Putative mRNA cleavage factor family protein | CFIm25 | 0\|0 | **2\|3** |
| Nucleic-acid binding |  |  |  |  |
| TGGT1_261960 | C2H2-type domain-containing protein | - | **9\|12** | 0\|0 |
| TGGT1_254220 | RRM domain-containing | - | 0\|0 | **6\|9** |
| Other locations & functions |  |  |  |  |
| TGGT1_240600 | Putative chaperonin cpn60 | - | **3\|3** | 0\|0 |
| TGGT1_269920 | Phosphatidylserine decarboxylase 2 proenzyme | - | **2\|2** | 0\|2 |

<sup>a</sup> protein that was immunoprecipitated
<sup>b</sup> bolded peptide counts are significantly enriched over parental (Padj ≤ 0.05; Student's T-test with Benjamini-Hochberg correction)
<sup>c</sup> manually included in list but did not meet statistical significance threshold due to high variability.

The YTH2 protein, which encodes an N-terminal YTH domain, also shares homology with CPSF30, though this is restricted to the YTH domain alone (Table 3). While fewer proteins were identified as interacting partners with YTH2^HA^, one included a core member of the 3’-end cleavage factor I complex and a second shares strong homology with the N-terminal region of mammalian Rb-binding protein 6 (RBBP6), which has been shown to play a role in 3’-end formation in yeast and mammals (34, 35). An additional interacting protein was an RRM-containing protein, which conceivably could play a role in binding mRNA during 3’-end formation.

Collectively, these findings indicate that both YTH m6A reader proteins in *Toxoplasma* are components of the 3’-end mRNA cleavage and polyadenylation machinery, and that m6A deposition within the YGCAUGCR consensus motif serves to facilitate pre-mRNA 3’-end formation.

## Discussion

### The *Toxoplasma* m6A writer complex possesses distinct features

In this study, we demonstrated that the *Toxoplasma* m6A writer complex consists of METTL3, METTL14, WTAP, a divergent VIRMA, and a writer-associated protein (WAP1) that remains to be characterized. Recently, affinity purification of the *P. falciparum* METTL3 homologue revealed that the writer complexes share a similar composition in both organisms, since METTL3, METTL14, WTAP, and the unannotated VIRMA (PF3D7_1366300) were co-purified (28). The *Toxoplasma* m6A writer complex also shares some features of that in higher eukaryotes, such as a labile interaction between the METTL3-METTL14 and the WTAP portions of the complex. Studies in *Drosophila melanogaster* have determined that the interaction between the METTL3-METTL14 heterodimer and the WTAP portion of the writer complex are sensitive to high salt concentrations such as those that we used during the nuclear extraction process (10). Alternatively, it has also been proposed that the METTL3-METT14 and WTAP portions of the complex may have distinct and non-overlapping functions (10).

Several reports have indicated that the catalytically active complex requires both METTL3 and METTL14 in a 1:1 stoichiometry (36, 37); it is therefore tempting to speculate that the lower amount of METTL14 in relation to METTL3 is biologically significant (Fig. 2). For example, METTL14 limitation may help regulate writer complex functionality in *Toxoplasma.* Whether METTL14 expression is regulated by cell cycle, environmental stimuli, or parasite differentiation are important areas of future investigation. Additionally, understanding the role of the novel WAP1 subunit in modulating m6A writer activity remains to be explored. Given that each member of the writer complex is a fitness-conferring gene for tachyzoite growth (16), depletion of METTL3 halts parasite growth (Fig. 4), and that the complex contains distinct members (Fig. 2D), it seems reasonable to conclude that deposition of the m6A mark is crucial to parasite viability and may be a good target for generating future anti-*Toxoplasma* therapies.

### m6A contributes to marking the 3’-end of newly synthesized transcripts

Since m6A deposition did not show stage-specific regulation (Fig. 3), and the parasite does not appear to encode m6A eraser proteins (Table 1), we propose that the m6A mark may be non-reversible and less dynamic than what has been proposed in higher eukaryotes (38, 39). Like *Toxoplasma, P. falciparum* also lacks identifiable m6A eraser proteins (40). Taken along with the essential requirement of the m6A mark, these characteristics may speak to its involvement in coordinating essential house-keeping functions in apicomplexans.

The m6A writer complex is known to modify mRNA co-transcriptionally (9). In other systems, this occurs on specific consensus sequences, supposedly due to recognition of these motifs by components of the writer complex. While animals, plants, and brewer’s yeast share similar motifs of DRACH, RRACH, and RGAC, respectively (8, 41,42), these motifs are varied in parasites such as *P. falciparum* (GGACA) (28) and *Trypanosoma brucei* (CAU) (29). Using an m6A-enrichment strategy paired with high-throughput RNAseq, we identified a strongly enriched YGCAUGCR motif with a centrally-located adenosine that represents the methylated residue (Fig. 4C). This motif is strongly enriched near the annotated 3’-boundaries of mRNAs, raising the prospect that m6A marks the 3’-end of pre-mRNAs.

In other species, pre-mRNAs are processed at their 3’-termini when a collection of sequence motifs are recognized by the cleavage and polyadenylation machinery. In animals, these are categorized as the upstream sequence element, the polyadenylation signal (PAS), the cleavage site, and a GU-rich downstream sequence element (43). The PAS consists of an AAUAAA motif located within 10-35nt upstream of the cleavage site, but this is only conserved in ~70% of transcripts (43). A similar organization exists in plants: the far upstream element (FUE) precedes an A-rich near upstream element (NUE) located 10-40nt upstream of the U-rich cleavage element (44, 45). Strict sequence conservation of the NUE is less important than what is seen in animals (45).

A recent attempt to identify the PAS of *Toxoplasma* and other related apicomplexan parasites was unable to define specific motifs (46), suggesting that, like plants, strict sequence conservation is not required. However, a distinct enrichment of adenosine residues was noted to be present ~20 nt upstream of the cleavage site (46) which could be supportive of a role for m6A in coordinating the process. We propose that the m6A motif, which is located in close proximity upstream of annotated 3’-boundary (Fig. 3E-F), operates analogously to the PAS/NUE in at least a subset of transcripts. Consistent with this idea, disruption of m6A deposition preferentially affects the steady state levels of *TGGT1_202650,* an mRNA that encodes an m6A motif near its 3’-end (Fig. 3E, 4C). We also identified an AU-rich motif that was enriched to within 10-40 nt downstream of the m6A element, putting it in a range that is consistent with it operating as an analogue to the U-rich cleavage element of plants. Specific mapping of the relationship between the *Toxoplasma* PAS/NUE and putative AU-rich cleavage element will be aided with increased assignment and improved boundary definition of 3’-UTRs genome-wide.

### The m6A system directs mRNA 3’-end formation in *Toxoplasma*

While the locale of the YGCAUGCR m6A motif may imply a role for the m6A system in 3’-end formation, the near complete purification of the cleavage and polyadenylation machinery along with the only two YTH domain-containing proteins, strongly positions the m6A system as a foundational piece driving this crucial process. The functions and individual constituents of the various cleavage and polyadenylation machinery are reviewed elsewhere (32, 43, 45); however, they can be broken down into cleavage and polyadenylation specificity factors (CPSF), cleavage stimulation factors (CSTF), and cleavage factors I and II (CFs).

CPSF30 plays a central role in coordinating the machinery associated with processing the 3’-end of transcripts. Interestingly, *Arabidopsis thaliana* produces two CPSF30 isoforms that arise from an alternative polyadenylation event (32, 33). The N-terminus of CPSF30 contains CCCH-type ZnF motifs that are responsible for binding the FUE (47) and is present in both CPSF30 isoforms whereas the longer version, CPSF30-L, is extended at its C-terminus and encodes an YTH domain, consistent with the protein architecture of apicomplexan YTH1 proteins (48). Plant CPSF30-L plays a role in nitrate signaling by mediating alternative polyadenylation events (49), and mediates the polyadenylation of tandemly duplicated genes in an m6A-dependent manner (50), suggesting functional conservation of the m6A-polyadenylation axis from apicomplexan parasites to plants.

Although both YTH1 and YTH2 share homology with CPSF30 in plants, it is likely that only YTH1 plays the role of plant CPSF30-L in *Toxoplasma* since it shares the same domain architecture; in contrast, YTH2 homology is restricted to the YTH domain and lacks the ZnFs characteristic of CPSF30s in all other organisms. In addition, unlike YTH2, YTH1 co-purified with all the other CPSF and CSTF factors that are identifiable in the *Toxoplasma* genome (Table 3), which is consistent with the role of CPSF30 as a central component of this machinery.

The exact role YTH2 plays in the 3’ end formation of transcripts remains to be resolved; however, it did associate with clearly identifiable members of the cleavage machinery (Table 3). Furthermore, a recent study has determined that the *P. falciparum* YTH2 homologue binds mRNAs near their 3’-ends (40), providing additional support for it playing a similar role in apicomplexan parasites. Future studies are required to determine the exact role YTH2 plays in 3’-end formation.

### Concluding remarks

Herein, we explored each identifiable component of the m6A system in *Toxoplasma.* In so doing, we uncovered the role that the m6A mark plays in coordinating the 3’-end formation of transcripts. Due to the similarity of the machinery and the organization of the motifs surrounding the cleavage site of transcripts with plants, we propose that the m6A system plays an evolutionarily conserved role in this process from plants to apicomplexan parasites. This raises the tantalizing possibility of leveraging the m6A system for future *anti-Toxoplasma* therapies, especially since 3’-end processing has previously been suggested to be an attractive drug target in *Toxoplasma* and *P. falciparum* (51, 52).

The discovery that the m6A system is involved in transcript maturation raises several possible avenues for future investigations. For instance, we have yet to understand whether m6A plays a role in regulating the alternative polyadenylation of transcripts, a wide-spread phenomenon that has been detected in *Toxoplasma* and other apicomplexans (46, 53, 54). Additionally, since we only identified reader proteins in the nucleus and many additional proteins outside of the YTH family have been shown to recognize the m6A mark in other organisms (8), whether the m6A mark directs other post-transcriptional regulatory events and the existence of cytoplasmic-localized readers remain open questions.

### Experimental Procedures

#### Cell strains and culture conditions

All parasites were cultured in human foreskin fibroblasts (ATCC: SCRC-1041) using standard procedures. Fibroblasts were initially grown in DMEM supplemented with 10% heat inactivated FBS until they reached confluency. The type II *Toxoplasma gondii* strain ME49 (ATCC: 50611) was used to assess m6A localization and for MeRIPseq. RH strain parasites in which the HXGPRT and Ku80 genes (RHΔΔ) have been removed (17, 18) were used for endogenous tagging of genes of writer and reader proteins. RHΔΔ engineered to express a copy of the TIR1 (31), gifted by Dr. David Sibley (Washington University), was used to generate the METTL3^HA-AID^ line. All tachyzoite cultures were maintained in DMEM supplemented with 1% heat inactivated FBS and 5% CO_2_. Bradyzoite differentiation was induced by incubating cultures in RPMI media supplemented with 5% heat inactivated FBS and buffered with 50 mM HEPES (pH 8.3) in an ambient CO_2_ incubator. Media for bradyzoite cultures was changed daily to maintain alkaline pH.

#### Endogenous tagging of genes

All writer proteins as well as YTH1 tagged lines were made by transfecting an appropriate linearized pLIC vector (18) into RHΔΔ parasites to generate C-terminal tagging by single homologous crossover. Briefly, for each gene, a sequence corresponding to approximately 1kb upstream of the annotated coding sequence stop codon was amplified from genomic DNA and cloned into the PacI site of the pLIC_HA vector. All primers are listed in Table S5. The plasmids were linearized using a unique restriction site within the gene-specific sequence and 5 μg were transfected into the parasites using a Nucleofector (Lonza Biosciences). Clones were selected for the DHFR cassette with 2 μM pyrimethamine and the correct integration was validated by PCR of the genomic DNA.

The YTH2^HA^ and METTL3^HA-AID^ lines were made by co-transfecting pCas9-GFP (55) and a PCR amplicon encoding the selected tag and selectable marker into parasites to generate C-terminal tagging by double homologous crossover. Briefly, the guide RNA cassette from pCas9-GFP was mutagenized by PCR to target cleavage near the stop codon of the gene of interest. A PCR amplicon was generated from a pLIC series vector to contain the HA (or HA-AID) epitope and the DHFR cassette along with an approximate 40nt overlap of upstream and downstream of the cut site. All primers are listed in Table S5. Once generated, 1 μg pCas9-GFP plasmid and 2 μg donor PCR amplicon were transfected into parasites using a Nucleofector (Lonza Biosciences). Clones were selected for the DHFR cassette with 2 μM pyrimethamine and the correct integration was validated by PCR of the genomic DNA.

#### Affinity purification of HA-tagged proteins

All affinity purifications were performed using intracellular tachyzoites freshly isolated from HFF cells. Two independent experiments were conducted separately for the writer proteins, whereas two biological samples were processed at the same time to generate data for the reader proteins. Each experiment also included parental strain parasites to control for non-specific interactions. Parasites were collected from infected fibroblasts by scraping the monolayers and passing them through a 23-gauge syringe and 5 μm pore filter. The parasites were washed twice with in PBS and then processed for subcellular fractionation. Affinity purification of the writer components were conducted from nuclear extracts whereas those conducted for the reader proteins were performed on whole cell lysate. Nuclear extracts were obtained by first incubating parasites for 5 minutes on ice in low salt lysis buffer (50 mM HEPES-NaOH pH 7.5, 10 mM NaCl, 0.1% NP-40, 20% glycerol), centrifugation to pellet nuclei and resuspending the pellet in high salt lysis buffer (50 mM HEPES-NaOH pH 7.5, 420 mM NaCl, 0.4% NP-40, 20% glycerol). Whole cell lysate was generated by incubating in lysis buffer (50 mM Tris pH 7.4, 150 mM NaCl, 1 mM MgCl_2_ 0.5% NP-40, 10% glycerol) and then passaging through a 23-gauge syringe. After clarifying the lysate in a refrigerated centrifuge at 21,000 xg in a tabletop centrifuge, the lysate was precleared on mouse IgG magnetic beads (Cell Signaling, 5873S) for 1 h while rotating at 4°C. The supernatant was transferred onto α-HA magnetic beads (Thermo Fisher, 88836) that were prewashed with lysis buffer and incubated overnight while rotating at 4°C. The beads were washed 3x with lysis buffer (with 0.05% NP-40) and 3x with PBS then submitted for protein discovery with the Proteomics Core Facility at the Indiana University School of Medicine.

Data was filtered to show proteins that were not detected in the parental strains and for statistically significance. Fisher’s exact test with Benjamini-Hochberg correction was used to determine significance of writer protein interactions and Student’s t-test Benjamini-Hochberg correction was performed to identify the significance of the reader protein interactome.

#### Immunofluorescence assays

HFFs were grown to confluency on glass coverslips. Parasites were added and grown under tachyzoite or bradyzoite-inducing conditions. After the allotted growth time, coverslips were fixed for 15 minutes with paraformaldehyde. Coverslips were permeabilized and blocked in 1X PBS supplemented with 3% BSA and 0.2% Triton X-100 for 30 minutes at room temperature. Endogenously tagged proteins were detected by incubating coverslips in blocking buffer supplemented with α-HA primary antibody (Roche, 11867423001, 1:1,000) for 1 h. Similarly, the m6A mark was detected with polyclonal α-m6A (Abcam, ab151230, 1:1,000). Coverslips were stained with appropriate secondary antibodies (Invitrogen, A11006, A11007, A11008, 1:2,000) and 2 μg/ml Hoescht or 1 μg/ml DAPI to reveal DNA, and 1:300 rhodamine-conjugated *Dolichos biflorus* agglutinin (Vector Laboratories, FL-1031) to reveal the bradyzoite cyst wall as appropriate.

#### Parasite viability assays

Parasite replication assays were performed by inoculating confluent HFF monolayers. The parasites were allowed to invade for 4 h, then the media was changed to include either 500 μM IAA (Sigma, I2886) or an equivalent volume of DMSO vehicle. The cultures were grown for 16 h before fixation. After staining, 100 randomly chosen vacuoles were counted for each condition in triplicate. Two independent experiments were performed.

For plaque assays, 5,000 parasites/well were inoculated into 12-well plates. After 24 h, the media was aspirated and wells were treated either with 500 μM IAA or an equivalent amount of DMSO vehicle. After an additional 24 h, the media was aspirated and wells were treated with either IAA or DMSO according to the designated treatment regimen. After an additional 4 days of growth, the plates were fixed in ice cold methanol and stained with crystal violet. Plaques were counted for each well which included 4 replicates of each condition. At least two independent experiments were conducted with similar results. Significance was assessed by Student’s t-test assuming unequal variance.

#### Western blots

To collect protein lysate, infected monolayers were rinsed twice with PBS and lysed in RIPA buffer (25 mM Tris pH 7.4, 150 mM NaCl, 0.1% SDS, 1% NP-40, 0.5% sodium deoxycholate) supplemented with a protease and phosphatase inhibitor cocktail (Sigma, PPC1010). Lysates were sonicated on ice and clarified by centrifugation at 21,000 xg for 10 min at 4°C. Samples were run on NuPAGE gels and transferred to nitrocellulose membranes. Blots were blocked for 30 min in a 5% non-fat milk TBST solution. Primary antibodies against the HA epitope (Roche, 11867423001, 1:1,000) and the SAG1 loading control (Thermo Fisher, MA5-18268, 1:1,000) were incubated overnight with the blot while rocking overnight at 4°C. Bands were detected by chemiluminescence using appropriate HRP-conjugated secondary antibodies (GE Healthcare, NA935V, NA931V, 1:5,000).

#### Dot blot

Total RNA was extracted from extracellular parasites with TRIzol (Ambion) as per the manufacturer. An aliquot of the RNA was run on an agarose gel to check for RNA integrity and for the absence of the 28S rRNA band characteristic of host cell contamination. Genomic DNA was isolated from the same aliquot of parasites with

DNeasy Blood and Tissue kit (Qiagen) as per the manufacturer. Indicated amounts of nucleic acids were spotted onto Hybond-N+ membrane (GE Healthcare) with a dot blot apparatus (BioRad). The nucleic acid was fixed to the membrane with a Stratagene crosslinker. After the membrane had dried, total nucleic acid stained with SYBR Gold (Invitrogen) for 5 min. Next, the membrane was blocked for 30 min in a 5% fat-free milk solution for 30 minutes followed by a 1 h incubation with α-m6A antibody (Abcam, ab151230, 1:1,000). The m6A mark was detected by chemiluminescence using an HRP-conjugated secondary antibody (GE Healthcare, NA934V, 1:5,000).

#### Quantitation of m6A levels

Total RNA was isolated from purified METTL3^HA-AID^ parasites that were treated with vehicle or 500 μM IAA as outlined above. The amount of m6A modification was assessed from 200 ng total RNA using the colorimetric EpiQuik m6A methylation quantification kit as per manufacturer’s instructions (EpiGentek, P-9005-96). The m6A levels were determined from four biological replicates and significance was assessed using Student’s t-test assuming unequal variances.

#### RT-PCR

Total RNA was isolated from intracellular parasites and was reverse transcribed with Superscript III (Invitrogen, 18080051) as per the manufacturer’s instructions using an anchored oligo-dT. PCR was conducted with Phusion polymerase as per the manufacturer’s instructions (NEB, 0530L). All primers are listed in Table S5. Three biological samples were tested for each condition and yielded similar results.

#### MeRIPseq and data analysis

For each sample, three T-175 flasks containing confluent HFFs were inoculated with ME49 parasites at an MOI of 10 and grown under tachyzoite conditions for 2 days. The media was aspirated and replaced to allow for tachyzoite or bradyzoite growth conditions for a further 24 h. Parasites were harvested by rinsing monolayers in PBS, scraping, syringe passage, and filtration as described for affinity purification experiments. Total RNA was isolated from the parasites and quality was assessed by gel electrophoresis. mRNA was enriched using Arraystar Seq-Star poly(A) isolation kit as per manufacturer’s instructions and subsequently fragmented by heating at 94°C in fragmentation buffer (10 mM Tris pH 7.0, 10 mM Zn^2+^). An aliquot of mRNA was saved for RNA sequencing as an input control for m6A RNA immunoprecipitation sequencing (MeRIPseq). Fragmented RNA was incubated with a polyclonal α-m6A antibody (Synaptic Systems, 202003) for 2 h at 4°C followed by an additional 2 h incubation with Dynabeads. Three washes were conducted with IP buffer (10 mM Tris pH 7.4, 150 mM NaCl, 0.1% NP-40) followed by two additional washes in wash buffer (10 mM Tris pH 7.4, 50 mM NaCl, 0.1% NP-40). The m6A-enriched mRNA was eluted in elution buffer (10 mM Tris pH 7.4, 1 mM EDTA, 0.05% SDS, 40U proteinase K) at 50°C for 30 minutes and subsequently phenol-chloroform extracted and ethanol precipitated. RNAseq libraries were constructed with KAPA Stranded mRNA-seq kit as per the manufacturer and sequencing was performed on an Illumina HiSeq 4000 system.

The annotated *Toxoplasma* ME49 strain genomic information was downloaded from ToxoDB v45 (56). Sequencing reads were depleted of tRNA and rRNA reads *in silico* using Bowtie (57). The remaining reads were aligned to the ME49 genome with HISAT2 (58). Differential expression was determined with DESEQ2 (59). Deduplicated sequencing reads were used to call m6A peaks using MeTPeak (25) and differential m6A marks were assessed using MeTDiff (26). Determination of the m6A motif was conducted with DREME (27) and its positional enrichment relative to the transcriptional stop site was determined with CentriMO (30). Default settings were used for all bioinformatic analyses. All datasets from this work have been deposited in the NCBI GEO database at https://www.ncbi.nlm.nih.gov/geo/query/acc.cgi (accession number GSE165067: **Reviewer token: cduncggyjpsrdat**).

## Acknowledgements

We thank Rashmita Basu and Leetah Senkpeil for technical assistance as well as Dr. Ron Wek, members of the Sullivan Lab, and the Biology of Intracellular Pathogens group at IUSM for helpful discussions.

## Supplemental Legends

**Figure S1.** Web logos of each m6A-like motif and AU-rich motif found in MeRIPseq data and annotated 3’UTRs. Significance (e-value) is also displayed.

**Table S1.** Complete dataset from writer complex immunoprecipitations. Data from two independent experiments are shown for parental, WTAP^HA^, METTL3^HA^, METTL14^HA^

**Table S2.** Complete differential expression analysis from RNAseq data. Normalized read counts, fold change and associated statistics are shown for both RNAseq and MeRIPseq analysis.

**Table S3.** Genomic locations of m6A peaks. Data for tachyzoites and bradyzoite-induced samples are shown as are those for differential m6A mark analysis.

**Table S4.** Complete dataset from m6A reader protein immunoprecipitations. Data are presented for parental, YTH1^HA^, and YTH2^HA^.

**Table S5.** List of primers used in this study. Primer combinations and the expected product sizes are indicated for tagged gene.

## References

1. Scallan E, Hoekstra RM, Angulo FJ, Tauxe RV, Widdowson MA, Roy SL, et al. Foodborne illness acquired in the United States--major pathogens. Emerg Infect Dis. 2011;17(1):7–15.

2. Hoffmann S, Batz MB, Morris JG, Jr. Annual cost of illness and quality-adjusted life year losses in the United States due to 14 foodborne pathogens. J Food Prot. 2012;75(7):1292–302.

3. Montoya JG, Liesenfeld O. Toxoplasmosis. Lancet. 2004;363(9425):1965–76.

4. Flegr J, Dama M. Does the prevalence of latent toxoplasmosis and frequency of Rhesus-negative subjects correlate with the nationwide rate of traffic accidents? Folia Parasitol (Praha). 2014;61(6):485–94.

5. Montazeri M, Mehrzadi S, Sharif M, Sarvi S, Tanzifi A, Aghayan SA, et al. Drug Resistance in Toxoplasma gondii. Front Microbiol. 2018;9:2587.

6. Lence T, Paolantoni C, Worpenberg L, Roignant JY. Mechanistic insights into m(6)A RNA enzymes. Biochim Biophys Acta Gene Regul Mech. 2019;1862(3):222–9.

7. Yue H, Nie X, Yan Z, Weining S. N6-methyladenosine regulatory machinery in plants: composition, function and evolution. Plant Biotechnol J. 2019;17(7):1194–208.

8. Berlivet S, Scutenaire J, Deragon JM, Bousquet-Antonelli C. Readers of the m(6)A epitranscriptomic code. Biochim Biophys Acta Gene Regul Mech. 2019;1862(3):329–42.

9. Tong J, Flavell RA, Li HB. RNA m(6)A modification and its function in diseases. Front Med. 2018;12(4):481–9.

10. Knuckles P, Lence T, Haussmann IU, Jacob D, Kreim N, Carl SH, et al. Zc3h13/Flacc is required for adenosine methylation by bridging the mRNA-binding factor Rbm15/Spenito to the m(6)A machinery component Wtap/Fl(2)d. Genes Dev. 2018;32(5-6):415–29.

11. Dominissini D, Moshitch-Moshkovitz S, Schwartz S, Salmon-Divon M, Ungar L, Osenberg S, et al. Topology of the human and mouse m6A RNA methylomes revealed by m6A-seq. Nature. 2012;485(7397):201–6.

12. Lesbirel S, Wilson SA. The m(6)A-methylase complex and mRNA export. Biochim Biophys Acta Gene Regul Mech. 2019;1862(3):319–28.

13. Meyer KD. m(6)A-mediated translation regulation. Biochim Biophys Acta Gene Regul Mech. 2019;1862(3):301–9.

14. Kierzek E, Kierzek R. The thermodynamic stability of RNA duplexes and hairpins containing N6-alkyladenosines and 2-methylthio-N6-alkyladenosines. Nucleic Acids Res. 2003;31(15):4472–80.

15. Liu N, Zhou KI, Parisien M, Dai Q, Diatchenko L, Pan T. N6-methyladenosine alters RNA structure to regulate binding of a low-complexity protein. Nucleic Acids Res. 2017;45(10):6051–63.

16. Sidik SM, Huet D, Ganesan SM, Huynh MH, Wang T, Nasamu AS, et al. A Genome-wide CRISPR Screen in Toxoplasma Identifies Essential Apicomplexan Genes. Cell. 2016;166(6):1423–35.e12.

17. Fox BA, Ristuccia JG, Gigley JP, Bzik DJ. Efficient gene replacements in Toxoplasma gondii strains deficient for nonhomologous end joining. Eukaryot Cell. 2009;8(4):520–9.

18. Huynh MH, Carruthers VB. Tagging of endogenous genes in a Toxoplasma gondii strain lacking Ku80. Eukaryot Cell. 2009;8(4):530–9.

19. Zimmermann L, Stephens A, Nam SZ, Rau D, Kübler J, Lozajic M, et al. A Completely Reimplemented MPI Bioinformatics Toolkit with a New HHpred Server at its Core. J Mol Biol. 2018;430(15):2237–43.

20. Yue Y, Liu J, Cui X, Cao J, Luo G, Zhang Z, et al. VIRMA mediates preferential m(6)A mRNA methylation in 3’UTR and near stop codon and associates with alternative polyadenylation. Cell Discov. 2018;4:10.

21. Meyer KD, Saletore Y, Zumbo P, Elemento O, Mason CE, Jaffrey SR. Comprehensive analysis of mRNA methylation reveals enrichment in 3’ UTRs and near stop codons. Cell. 2012;149(7):1635–46.

22. Soête M, Camus D, Dubremetz JF. Experimental induction of bradyzoite-specific antigen expression and cyst formation by the RH strain of Toxoplasma gondii in vitro. Exp Parasitol. 1994;78(4):361–70.

23. Kim K. The Epigenome, Cell Cycle, and Development in Toxoplasma. Annu Rev Microbiol. 2018;72:479–99.

24. Hong DP, Radke JB, White MW. Opposing Transcriptional Mechanisms Regulate Toxoplasma Development. mSphere. 2017;2(1).

25. Cui X, Meng J, Zhang S, Chen Y, Huang Y. A novel algorithm for calling mRNA m6A peaks by modeling biological variances in MeRIP-seq data. Bioinformatics. 2016;32(12):i378–i85.

26. Cui X, Zhang L, Meng J, Rao MK, Chen Y, Huang Y. MeTDiff: A Novel Differential RNA Methylation Analysis for MeRIP-Seq Data. IEEE/ACM Trans Comput Biol Bioinform. 2018;15(2):526–34.

27. Bailey TL. DREME: motif discovery in transcription factor ChIP-seq data. Bioinformatics. 2011;27(12):1653–9.

28. Baumgarten S, Bryant JM, Sinha A, Reyser T, Preiser PR, Dedon PC, et al. Transcriptome-wide dynamics of extensive m(6)A mRNA methylation during Plasmodium falciparum blood-stage development. Nat Microbiol. 2019;4(12):2246–59.

29. Liu L, Zeng S, Jiang H, Zhang Y, Guo X, Wang Y. Differential m6A methylomes between two major life stages allows potential regulations in Trypanosoma brucei. Biochem Biophys Res Commun. 2019;508(4):1286–90.

30. Bailey TL, Machanick P. Inferring direct DNA binding from ChIP-seq. Nucleic Acids Res. 2012;40(17):e128.

31. Brown KM, Long S, Sibley LD. Plasma Membrane Association by N-Acylation Governs PKG Function in Toxoplasma gondii. mBio. 2017;8(3).

32. Hunt AG, Xing D, Li QQ. Plant polyadenylation factors: conservation and variety in the polyadenylation complex in plants. BMC Genomics. 2012; 13:641.

33. Chakrabarti M, Hunt AG. CPSF30 at the Interface of Alternative Polyadenylation and Cellular Signaling in Plants. Biomolecules. 2015;5(2):1151–68.

34. Di Giammartino DC, Li W, Ogami K, Yashinskie JJ, Hoque M, Tian B, et al. RBBP6 isoforms regulate the human polyadenylation machinery and modulate expression of mRNAs with AU-rich 3’ UTRs. Genes Dev. 2014;28(20):2248–60.

35. Vo LT, Minet M, Schmitter JM, Lacroute F, Wyers F. Mpe1, a zinc knuckle protein, is an essential component of yeast cleavage and polyadenylation factor required for the cleavage and polyadenylation of mRNA. Mol Cell Biol. 2001;21(24):8346–56.

36. Wang P, Doxtader KA, Nam Y. Structural Basis for Cooperative Function of Mettl3 and Mettl14 Methyltransferases. Mol Cell. 2016;63(2):306–17.

37. Liu J, Yue Y, Han D, Wang X, Fu Y, Zhang L, et al. A METTL3-METTL14 complex mediates mammalian nuclear RNA N6-adenosine methylation. Nat Chem Biol. 2014;10(2):93–5.

38. Jia G, Fu Y, Zhao X, Dai Q, Zheng G, Yang Y, et al. N6-methyladenosine in nuclear RNA is a major substrate of the obesity-associated FTO. Nat Chem Biol. 2011;7(12):885–7.

39. Zhou J, Wan J, Gao X, Zhang X, Jaffrey SR, Qian SB. Dynamic m(6)A mRNA methylation directs translational control of heat shock response. Nature. 2015;526(7574):591–4.

40. Govindaraju G, Kadumuri RV, Sethumadhavan DV, Jabeena CA, Chavali S, Rajavelu A. N(6)-Adenosine methylation on mRNA is recognized by YTH2 domain protein of human malaria parasite Plasmodium falciparum. Epigenetics Chromatin. 2020;13(1):33.

41. Shen L, Liang Z, Gu X, Chen Y, Teo ZW, Hou X, et al. N(6)-Methyladenosine RNA Modification Regulates Shoot Stem Cell Fate in Arabidopsis. Dev Cell. 2016;38(2):186–200.

42. Schwartz S, Agarwala SD, Mumbach MR, Jovanovic M, Mertins P, Shishkin A, et al. High-resolution mapping reveals a conserved, widespread, dynamic mRNA methylation program in yeast meiosis. Cell. 2013;155(6):1409–21.

43. Nourse J, Spada S, Danckwardt S. Emerging Roles of RNA 3’-end Cleavage and Polyadenylation in Pathogenesis, Diagnosis and Therapy of Human Disorders. Biomolecules. 2020;10(6).

44. Hunt AG. Messenger RNA 3’ end formation in plants. Curr Top Microbiol Immunol. 2008;326:151–77.

45. Bernardes WS, Menossi M. Plant 3’ Regulatory Regions From mRNA-Encoding Genes and Their Uses to Modulate Expression. Front Plant Sci. 2020;11:1252.

46. Stevens AT, Howe DK, Hunt AG. Characterization of mRNA polyadenylation in the apicomplexa. PLoS One. 2018;13(8):e0203317.

47. Addepalli B, Hunt AG. A novel endonuclease activity associated with the Arabidopsis ortholog of the 30-kDa subunit of cleavage and polyadenylation specificity factor. Nucleic Acids Res. 2007;35(13):4453–63.

48. Thomas PE, Wu X, Liu M, Gaffney B, Ji G, Li QQ, et al. Genome-wide control of polyadenylation site choice by CPSF30 in Arabidopsis. Plant Cell. 2012;24(11):4376–88.

49. Li Z, Wang R, Gao Y, Wang C, Zhao L, Xu N, et al. The Arabidopsis CPSF30-L gene plays an essential role in nitrate signaling and regulates the nitrate transceptor gene NRT1.1. New Phytol. 2017;216(4):1205–22.

50. Pontier D, Picart C, El Baidouri M, Roudier F, Xu T, Lahmy S, et al. The m(6)A pathway protects the transcriptome integrity by restricting RNA chimera formation in plants. Life Sci Alliance. 2019;2(3).

51. Sonoiki E, Ng CL, Lee MC, Guo D, Zhang YK, Zhou Y, et al. A potent antimalarial benzoxaborole targets a Plasmodium falciparum cleavage and polyadenylation specificity factor homologue. Nat Commun. 2017;8:14574.

52. Palencia A, Bougdour A, Brenier-Pinchart MP, Touquet B, Bertini RL, Sensi C, et al. Targeting Toxoplasma gondii CPSF3 as a new approach to control toxoplasmosis. EMBO Mol Med. 2017;9(3):385–94.

53. Wan KL, Chang TL, Ajioka JW. Molecular characterization of tgd057, a novel gene from Toxoplasma gondii. J Biochem Mol Biol. 2004;37(4):474–9.

54. Matrajt M, Platt CD, Sagar AD, Lindsay A, Moulton C, Roos DS. Transcript initiation, polyadenylation, and functional promoter mapping for the dihydrofolate reductase-thymidylate synthase gene of Toxoplasma gondii. Mol Biochem Parasitol. 2004;137(2):229–38.

55. Shen B, Brown KM, Lee TD, Sibley LD. Efficient gene disruption in diverse strains of Toxoplasma gondii using CRISPR/CAS9. mBio. 2014;5(3):e01114–14.

56. Harb OS, Roos DS. ToxoDB: Functional Genomics Resource for Toxoplasma and Related Organisms. Methods Mol Biol. 2020;2071:27–47.

57. Langmead B, Trapnell C, Pop M, Salzberg SL. Ultrafast and memory-efficient alignment of short DNA sequences to the human genome. Genome Biol. 2009;10(3):R25.

58. Kim D, Langmead B, Salzberg SL. HISAT: a fast spliced aligner with low memory requirements. Nat Methods. 2015;12(4):357–60.

59. Love MI, Huber W, Anders S. Moderated estimation of fold change and dispersion for RNA-seq data with DESeq2. Genome Biol. 2014;15(12):550.

